# DNA fragment length analysis using machine learning assisted vibrational spectroscopy

**DOI:** 10.64898/2026.02.27.708538

**Authors:** Rashad Fatayer, Waseem Ahmed, Irene Szeto, Stephen-John Sammut, Ganapathy Senthil Murugan

## Abstract

DNA length analysis is essential for genomic workflows including next-generation sequencing and fragmentomics based diagnostics. Conventional approaches typically require large, expensive instrumentation and sample-destructive protocols with long processing times. Here we present a rapid, label-free approach integrating vibrational spectroscopy with deep learning to quantify DNA fragment length distributions. We demonstrate that ATR-FTIR and Raman spectroscopy capture length-dependent spectral features arising from phosphate backbone, nucleobase, and structural vibrations. Machine learning models trained on spectra acquired from purified monodisperse DNA (50-300 bp) predicted DNA length with high accuracy (R^2^=0.92-0.94), with multimodal fusion improving performance to R^2^=0.96. A convolutional neural network trained on 35 DNA mixtures comprising molecules of different lengths also successfully deconvoluted their fragment length profile. Transfer learning enabled adaptation to biological samples, achieving low prediction error (RMSE=0.3-7.2%, Δμ=12 bp). Importantly, the method requires only 4 μL sample and 15 minutes passive drying, with no consumables beyond cleaning materials, and allows full sample recovery. This establishes vibrational spectroscopy as a scalable alternative for DNA length quantification.

---

The quantification of DNA fragment length is a critical step in many genomic workflows, including next generation sequencing library preparation ^1,2^. Beyond sequencing, DNA fragment length distribution in biofluids such as blood is emerging as a key biomarker in several diagnostic contexts ^3^. For example, in cancer, DNA released by tumour cells into the circulation is shorter (90-150 base pairs) than that released by normal cells (∼167 base pairs) ^4^, and this difference can be detected by sequencing. Similarly, DNA specific length profiles allow for the differentiation of foetal DNA from maternal DNA in prenatal screening ^5^, and the identification of viral DNA in infectious disease monitoring^6^.

Conventional methods for fragment length analysis include gel electrophoresis, which estimates length based on DNA migration through a gel matrix ^7^, but offers limited resolution and requires lengthy, multi-step workflows. Higher resolution technologies such as automated capillary electrophoresis and sequencing offer higher accuracy and precision but are more expensive, require larger instrumentation and involve more stringent sample preparation protocols ^8–10^. These constraints limit the accessibility of high-resolution DNA fragment length quantification, particularly in laboratories operating under budgetary limitations. To address this, we explored an alternative approach that bridges the gap between accessible, low-resolution methods and high-cost, high-precision platforms. We designed and evaluated a novel methodology based on vibrational spectroscopy, specifically Attenuated Total Reflectance-Fourier Transform Infrared (ATR-FTIR) and Raman spectroscopies, combined with machine learning to enable label-free, non-destructive DNA fragment length quantification. This technology probes the vibrational modes of chemical bonds, producing spectral features that reflect the underlying molecular structure ^11,12^.

In this proof-of-principle study (Fig. 1), we present a novel method to investigate whether length dependent structural features in DNA can be detected using vibrational spectroscopy. We show that variations in band intensities and wavenumber positions carry fragment length information that allows accurate quantification of DNA fragment lengths. Furthermore, we demonstrate that a 1D convolutional neural network (1D-CNN) can estimate fragment length proportions in mixtures from ATR-FTIR spectra with low error. Collectively, these findings suggest that machine learning assisted vibrational spectroscopy may provide a rapid and low-cost option for fragment length analysis, particularly in settings where cost-effective, fast and accurate workflows are required.

**Fig.1:**
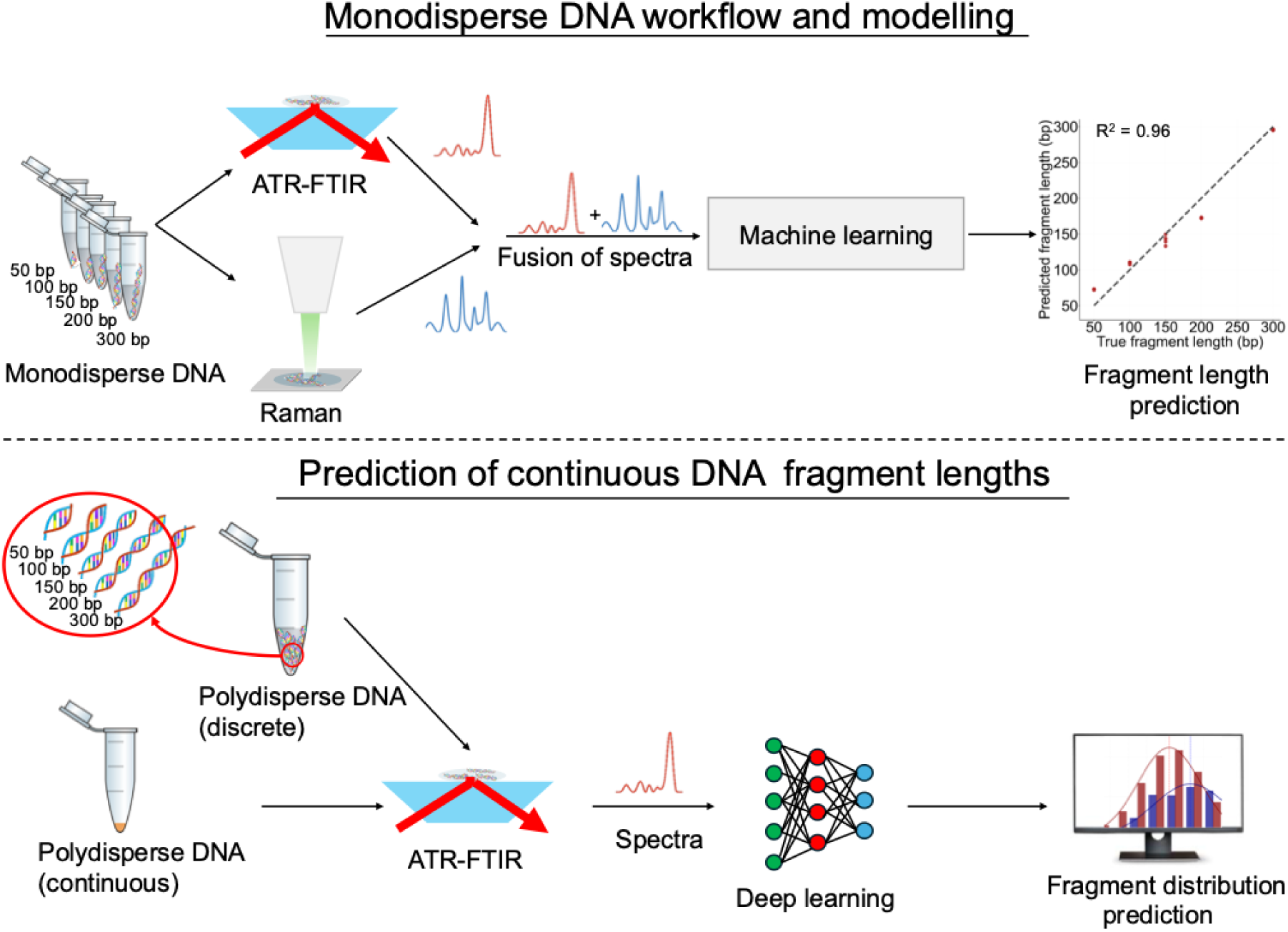
Study overview. Monodisperse DNA samples containing single fragment lengths were analysed using ATR-FTIR and Raman spectroscopy. Spectra were combined using low-level data fusion and used as input to a Partial Least Squares Regression (PLSR) model for DNA fragment length prediction. For polydisperse DNA samples, two types of sample types were prepared. The first comprised defined mixtures of monodisperse DNA fragments combined in proportions to generate discrete polydisperse DNA mixtures. The second consisted of polydisperse DNA produced by shearing of genomic DNA, producing a continuous fragment length distribution. Discrete polydisperse samples were analysed using ATR-FTIR spectroscopy, and the resulting spectra were used to train a one-dimensional convolutional neural network (1D-CNN). Transfer learning approach was used (from discrete polydispersed to continuous polydispersed) to quantify DNA fragment length distribution in biological samples.

## DNA fragment length in monodisperse solutions

To establish proof of principle, we first investigated whether vibrational spectroscopy could discriminate between monodisperse DNA solutions comprising a single fragment length (measured in base pairs, bp). Solutions containing DNA fragments of five discrete lengths (50, 100, 150, 200, and 300 bp), each at an identical concentration, were analysed using ATR-FTIR spectroscopy (Methods). Following spectral averaging (Figure 2a), fragment length-dependent variations in both absorbance intensity and peak position were observed across the full measured range (750-4000cm^-1^).

**Fig. 2:**
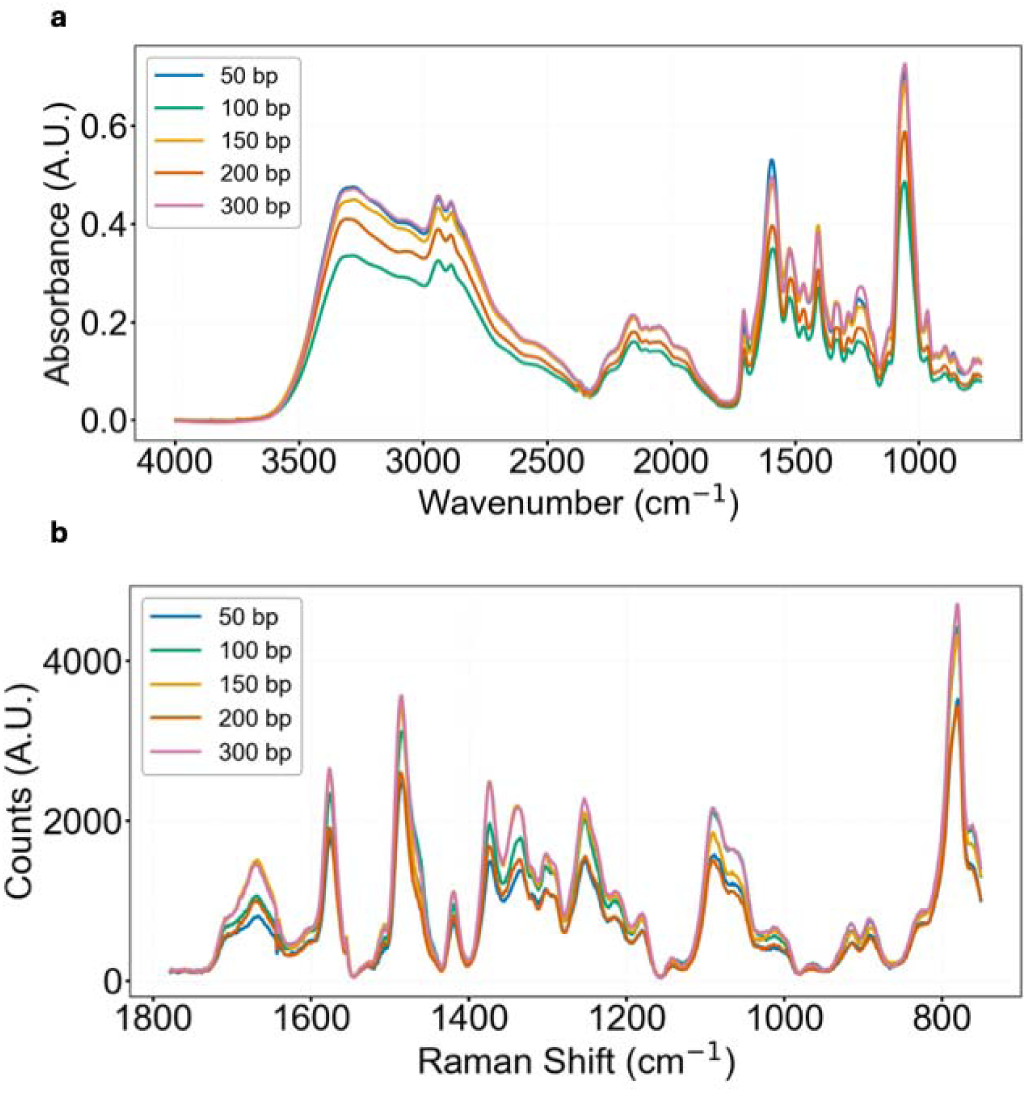
Spectroscopic analysis of monodisperse DNA solutions containing different fragment lengths. **(a)** Averaged FTIR and **(b)** averaged Raman spectra of DNA fragments (50-300 bp).

The most prominent DNA associated features were observed in (1) the symmetric PO_2_ ^-^ stretching band (∼1080 cm^-1^), which showed the largest change in intensity across fragment lengths, (2) the asymmetric PO_2_ ^-^ stretching region (∼1220-1250 cm^-1^), and (3) nucleobase associated vibrations from C=O, C=N, and C=C bonds (∼1500-1700 cm^-1^) ^13–16^. Within the ∼1250 cm^-1^ band, shorter fragments displayed a shift towards higher wavenumbers. This behaviour is consistent with the structural organisation of DNA, as these spectroscopic bands originate from sugar phosphate backbone vibrations. Fragment length-dependent differences were also observed within the nucleobase region (∼1500-1700 cm^-1^), likely reflecting base stacking interactions ^17,18^ associated with increasing structure regularity in longer fragments. Variations between fragment lengths were also detected around ∼3300 cm^-^¹ within the broad N-H and O-H stretching regions consistent with changes in hydrogen bonding associated with DNA hydration and base groups ^19^.

We subsequently analysed these DNA monodisperse solutions using Raman spectroscopy (Figure 2b). The Raman spectra showed that the strongest band observed was at ∼785 cm^-1^, which is associated with O-P-O stretching vibrations, together with cytosine and thymine ring breathing modes ^20^. Strong contributions were also observed in the 1300-1800 cm^-^¹ region typically associated with nucleobase vibrations ^21,22^. A further prominent feature occurred at ∼1085 cm^-1^, corresponding to symmetric PO_2_ ^-^ stretching ^23^, consistent with the ATR-FTIR findings where this band was also identified as having strong fragment length dependence. Additional bands were observed between 1250 cm^-1^ and 1375 cm^-1^ originating from cytosine, thymine, and adenine vibrations, as well as purine ring stretching modes. Other regions seen from 1485cm^-1^ to 1580 cm^-1^ are assigned to adenine, guanine, and cytosine ring stretching modes, with a sharp band at ∼1660 cm^-1^ reflecting C=O stretching of thymine and guanine ^21^.

Finally, across all DNA with various fragment lengths, the Raman spectral features remained consistent in overall shape but showed length dependent intensity variations, most notably in the ∼1085 cm^-1^ and ∼1450 cm^-1^ bands. Wavenumber shifts were also observed relative to the ∼1250 cm^-1^ nucleobase associated region, however this effect was not as significant as observed in ATR-FTIR.

In summary, we show that both ATR-FTIR and Raman spectra display DNA length-dependent spectral changes in monodisperse solutions containing DNA fragments of identical lengths, with differences arising from backbone, nucleobase and hydrogen bonding related vibrations.

## Multivariate analysis of monodisperse DNA fragment length solutions

To explore whether the spectral features identified by ATR-FTIR and Raman were sufficient to predict DNA fragment length in monodisperse solutions, we first split the data into training and test sets and applied Partial Least Squares Discriminatory Analysis (PLS-DA) to each modality separately. PLS-DA was used to incorporate the fragment length labels and extract latent variables that maximise class separation based on the underlying spectral features. Five PLS latent variables were retained for the PLS-DA models. For both ATR-FTIR and Raman datasets, the second and third latent variables provided the strongest class separation.

Across both modalities, latent space projections showed clustering of the five DNA length groups (Figures 3a and 3b). In ATR-FTIR data, the dominant loadings (Supplementary Fig. 1) mapped primarily to the 1000-1800 cm^-1^ region, consistent with phosphate backbone and nucleobase associated vibrations. In Raman spectra, the loadings (Supplementary Fig. 2) focused on shorter wavenumbers ∼670-870 cm^-1^, consistent with ring breathing modes from DNA bases ^21^. Additional discriminative features were observed around ∼1575 cm^-1^ which are also attributable to base ring breathing vibrations.

**Fig. 3:**
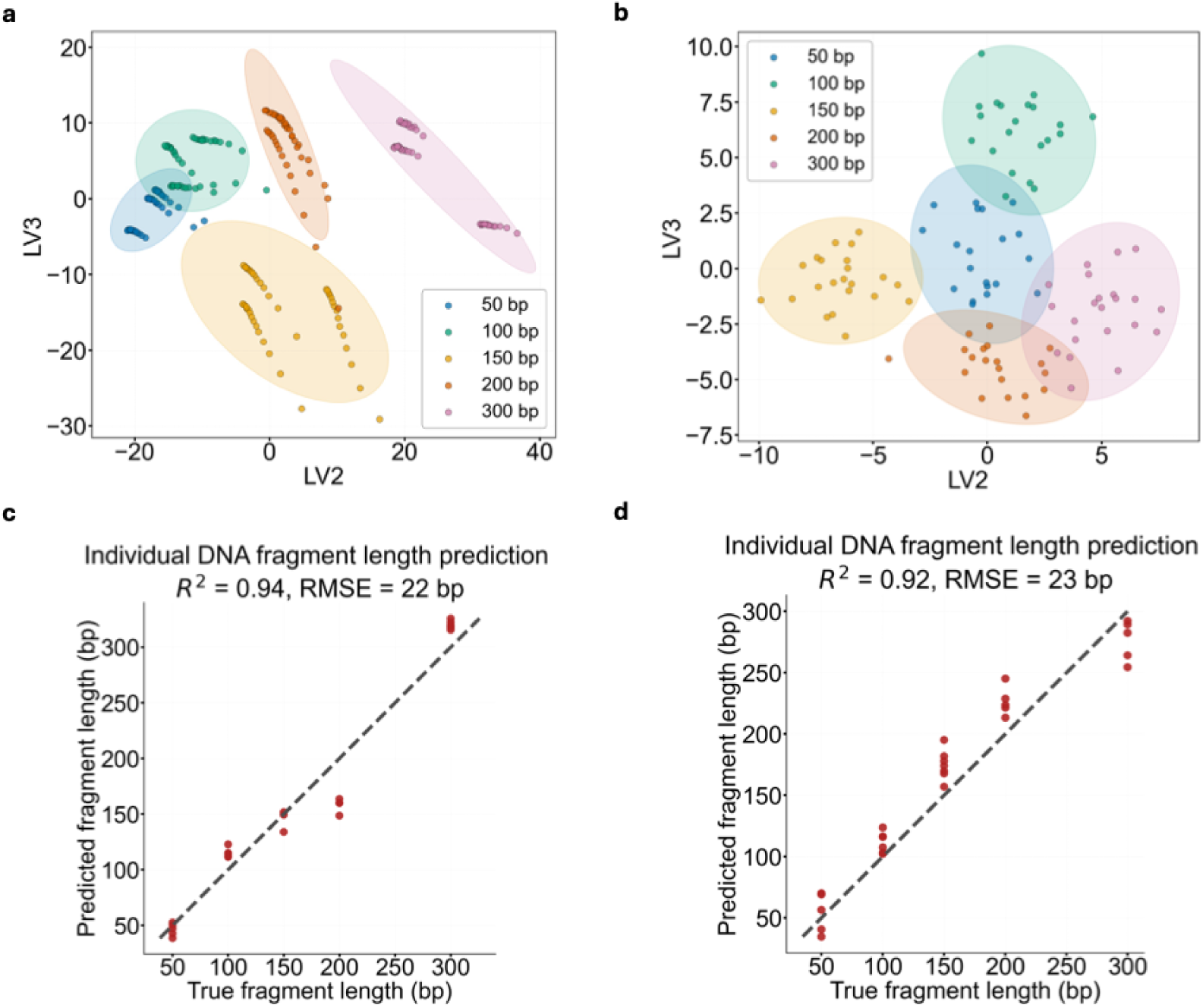
Spectroscopic and machine learning analysis of individual DNA fragment length. Partial Least Squares (PLS) scores plot of **(a)** FTIR and **(b)** Raman data. Solutions with different fragment lengths cluster based on their scattering features. Scatter plots showing the performance of PLSR models trained on **(c)** FTIR and **(d)** Raman data. Ellipses in (**a) and (b)** are representing the 95% confidence interval for each group.

Following this analysis, we developed Partial Least Squares Regression (PLSR) models to predict DNA fragment length directly from the spectra. To ensure that the models focused on all relevant biochemical information, the analysis was restricted to the strongest DNA specific vibrational windows (750-1800 cm^-1^ for FTIR and 600-1800 cm^-1^ for Raman). Data splitting in the PLSR models followed the experimental structure to prevent data leakage: for ATR-FTIR, samples measured were split by replicate (first two for training and cross-validation, third for testing), while Raman data were split by physical laser spot position (see Methods for more details). Models were optimized by selecting the number of latent variables based on cross-validation performance, with Van der Voet’s F-test applied to prevent overfitting (Supplementary Fig. 3 and 4, and Methods).

Both PLSR models trained independently using ATR-FTIR and Raman spectra achieved comparable performance on their respective test sets. For FTIR, the model achieved an R^2^ of 0.94 with an RMSE of 22 bp (Figure 3c), while the Raman model yielded an R^2^ of 0.92 and an RMSE of 23 bp (Figure 3d).

In summary, these results show that statistical modelling of ATR-FTIR and Raman spectra enables accurate length quantification of monodisperse DNA solutions containing a single length.

## Data fusion enhances fragment length prediction

We next integrated ATR-FTIR and Raman spectra using data fusion (Methods). Although both techniques detect molecular vibrations, ATR-FTIR measures dipole moment changes while Raman spectroscopy measures polarizability changes. As a result, when the techniques are used together they have been shown to be complementary improving predictive performance ^24–26^. For example, while the observed symmetric PO ^-^ stretch (∼1080 cm^-1^) was a dominant, length sensitive feature in FTIR, this was weaker in the Raman spectrum. Conversely, the O-P-O associated backbone vibration (∼785 cm^-1^) was a dominant peak in Raman but was almost absent in FTIR.

To fully evaluate whether integrating data from both modalities improved model fit, we first created a fused dataset using low level feature concatenation. For each sample, the Raman spectrum (originally 600-1800 cm^-1^) was appended to the end of the FTIR spectrum (originally 750-1800 cm^-1^) to create a single, unified data vector (Figure 4a). Applying the PLSR methodology developed for the independent modalities to this fused dataset improved the model’s predictive ability. By combining spectral data from ATR-FTIR and Raman spectroscopies, the data fusion model achieved an R^2^ of 0.96 and reduced the RMSE to 17 bp (Figure 4b), outperforming the FTIR only model (RMSE 22 bp) and the Raman only model (RMSE 23 bp).

**Fig. 4:**
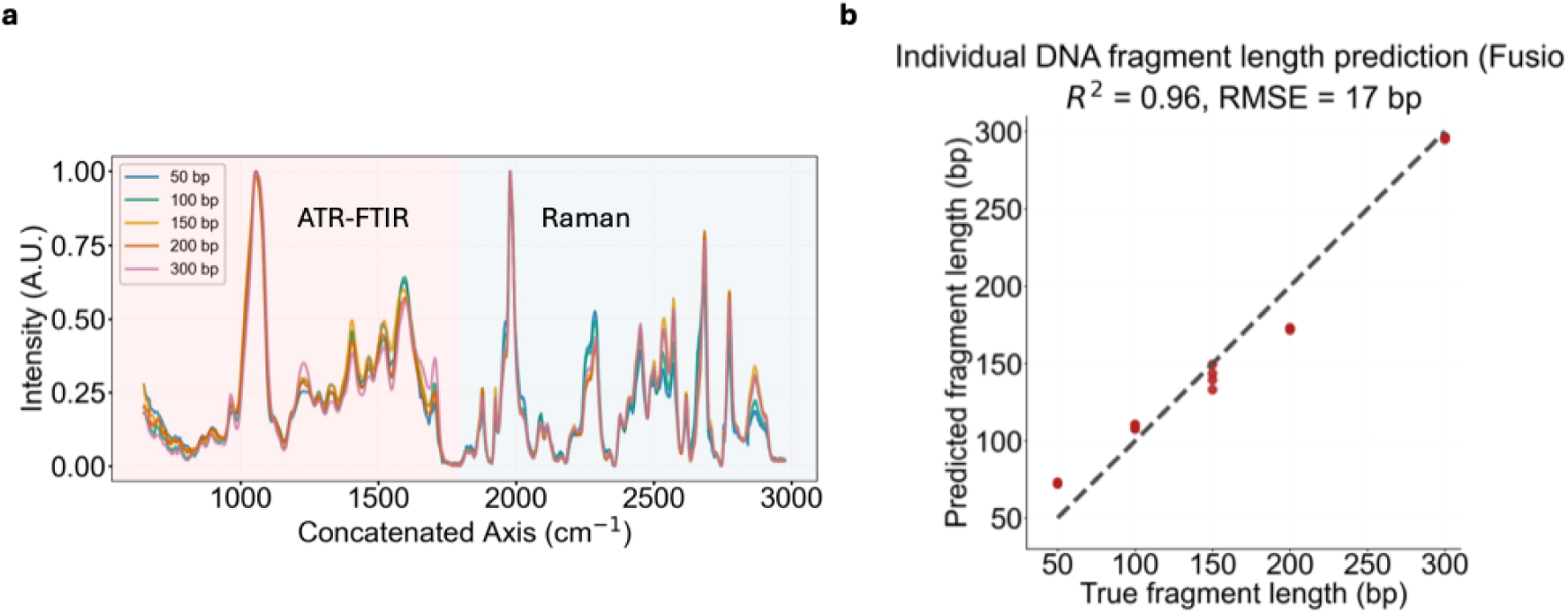
Data fusion of FTIR and Raman spectra increases predictive accuracy. **(a)** Visualization of low level feature concatenation. The x axis represents a synthetic axis created for continuity, showing the FTIR spectrum (up to ∼1800 cm^-1^) and the Raman spectrum (from ∼1800 cm^-1^ onwards). **(b)** Regression plot for a PLSR model trained on the fused dataset.

This work represents the first application of multimodal vibrational spectroscopy for quantitative DNA fragment length analysis, introducing a methodology that combines complementary ATR-FTIR and Raman data to improve prediction accuracy in monodisperse DNA solutions.

## Developing a 1D-CNN model on polydisperse mixtures with discrete DNA fragment lengths

We next sought to determine whether fragment length information could be deconvoluted from polydisperse DNA solutions containing discrete DNA fragment lengths. We generated 35 polydisperse DNA mixtures comprising all five fragment lengths (50 bp, 100 bp, 150 bp, 200 bp, and 300 bp) in systematically varying proportions. The sample set included monodisperse samples, binary, ternary, quaternary, and quintenary mixtures (Figure 5a and Methods). ATR-FTIR spectra were collected for all 35 mixture compositions, providing a comprehensive dataset to train and validate the fragment length model. Such mixtures produce complex, overlapping spectral features not easily resolved by linear models (Supplementary Fig. 5 and 6).

**Fig. 5:**
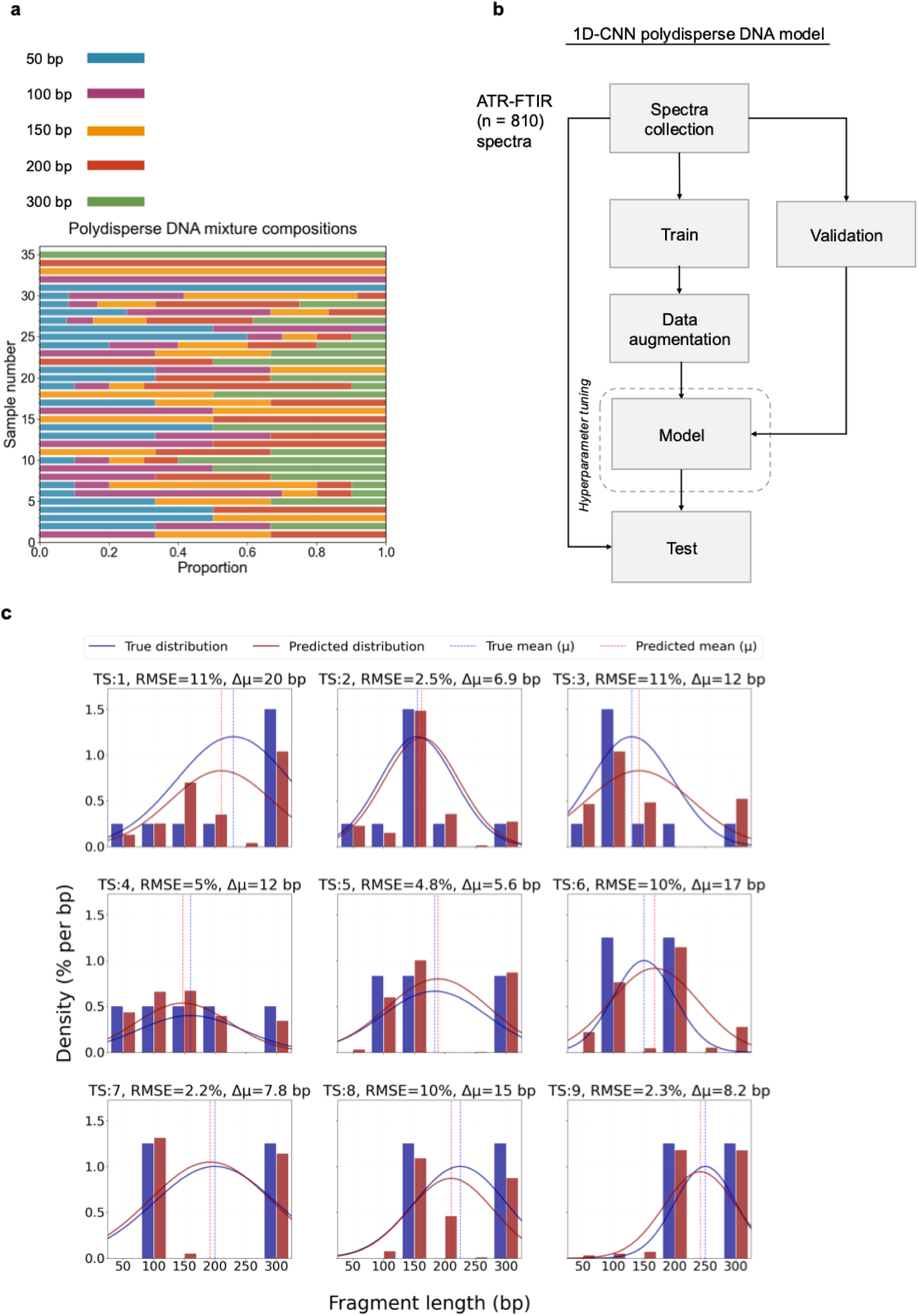
1D-CNN workflow and performance on polydisperse DNA mixtures containing discrete fragment lengths. **(a)** Stacked bar chart showing the composition of 35 polydisperse DNA mixture samples generated by mixing DNA solutions containing fragments of defined lengths (50 bp, 100 bp, 150 bp, 200 bp, and 300 bp). Each bar represents a unique mixture composition, with colours indicating the proportion of each fragment length present. (**b)** Machine learning workflow for model development. **(c)** Predicted fragment length distributions produced by the 1D-CNN model (red bars), with the corresponding Gaussian estimate overlaid as a solid red line. True fragment length probabilities, derived from the known mixture compositions, are shown as blue bars with their corresponding Gaussian estimates indicated by solid blue lines. Results are shown for nine representative samples from the independent test set (TS). Vertical dotted lines indicate the weighted mean fragment length for both predicted (red) and true (blue) distributions.

To address this, we developed a 1D-CNN (Figure 5b). This deep learning architecture is optimised for identifying local patterns in spectral data and capable of disentangling spectral overlap ^27–30^. The 1D-CNN was trained on the measured ATR-FTIR spectra from the DNA mixtures for simultaneous prediction of all fragment percentages. Following a similar train and test data partitioning method as the PLSR models, the first two replicate spectra measurements were used for training, with 20% of the training reserved for validation, while the third replicate set served as an independent test. To improve generalisation and reduce overfitting, the training set was augmented six-fold using intensity scaling, baseline shifts, and the addition of low amplitude noise (Supplementary Fig. 7). Model optimisation was performed by minimising the loss (Supplementary Fig. 8) between the predicted and true fragment proportions on the training set, with validation loss used to monitor convergence and generalisation.

As the model outputs discrete proportions for each fragment class, we further represented the predicted mixtures as continuous Gaussian distributions using their weighted mean and standard deviation. These Gaussian representations were used for visualisation and comparison, and not as a constraint during training. This smoothing step provided a more intuitive basis for comparing fragmentation patterns between samples, particularly when assessing mixtures enriched in shorter versus longer fragments where fragmentation profiles can serve as diagnostic biomarkers ^4,31,32^.

The 1D-CNN achieved accurate DNA length proportion estimation across the entire test set (Supplementary Fig. 9 and 10). The lowest error was for 50 bp fragments (RMSE = 4.7%) and the highest for 300 bp fragments (RMSE = 9.0%). Across mixtures, the model achieved an average RMSE of 6.5% and a mean distribution difference (Δμ) of 12 bp between true and predicted profiles (Figure 5c).

In summary, these results demonstrate that a 1D-CNN trained on augmented spectral data can resolve vibrational signatures from mixtures of DNA with varying fragment lengths and accurately reconstruct the underlying length distributions. This represents the first application of machine learning to vibrational spectroscopy for DNA fragmentomic profiling, providing a rapid and label-free alternative to separation-based methods for analyzing complex biological samples.

## Quantifying complex DNA mixtures with continuous fragment lengths

We next explored whether our methodology could be extended to estimate fragment length distributions in biological DNA samples comprising continuous, rather than discrete, fragment lengths. To do this, we prepared 11 sheared DNA samples spanning a range of fragment length distributions and quantified their fragment length distribution using gel electrophoresis, the current gold standard (Methods). Gel electrophoresis confirmed that all samples contained broad and continuous fragment length distributions rather than discrete populations. Mean fragment lengths ranged from ∼195 to ∼240 bp, while median lengths spanned from ∼175 bp to ∼235 bp. Quantitative binning of the electropherograms revealed overlap across neighbouring fragment lengths for all samples, with the most occurring fragment typically centred within the 150-200 bp and 200-250 bp bins (Supplementary Table 4. and 5.). In general, shorter fragments (<100 bp) were consistently present at low abundance while longer fragments (250-350 bp) increased progressively across the sample series.

We subsequently acquired the ATR-FTIR spectra for the samples. Given the limited number of biological samples, the collected spectral dataset was divided into six samples for model training and five for testing, with both subsets harbouring comparable DNA fragment length distributions. We employed a transfer learning strategy rather than training a new model from scratch (Figure 6a). Specifically, the 1D-CNN model previously trained on the 35 purified mixtures was repurposed as a feature extractor, and its final layers were fine-tuned using the biological training set. This two-stage approach enabled the model to adapt to the domain shift between purified and biological DNA spectra while retaining the spectral features learned from the larger original dataset.

**Fig.6:**
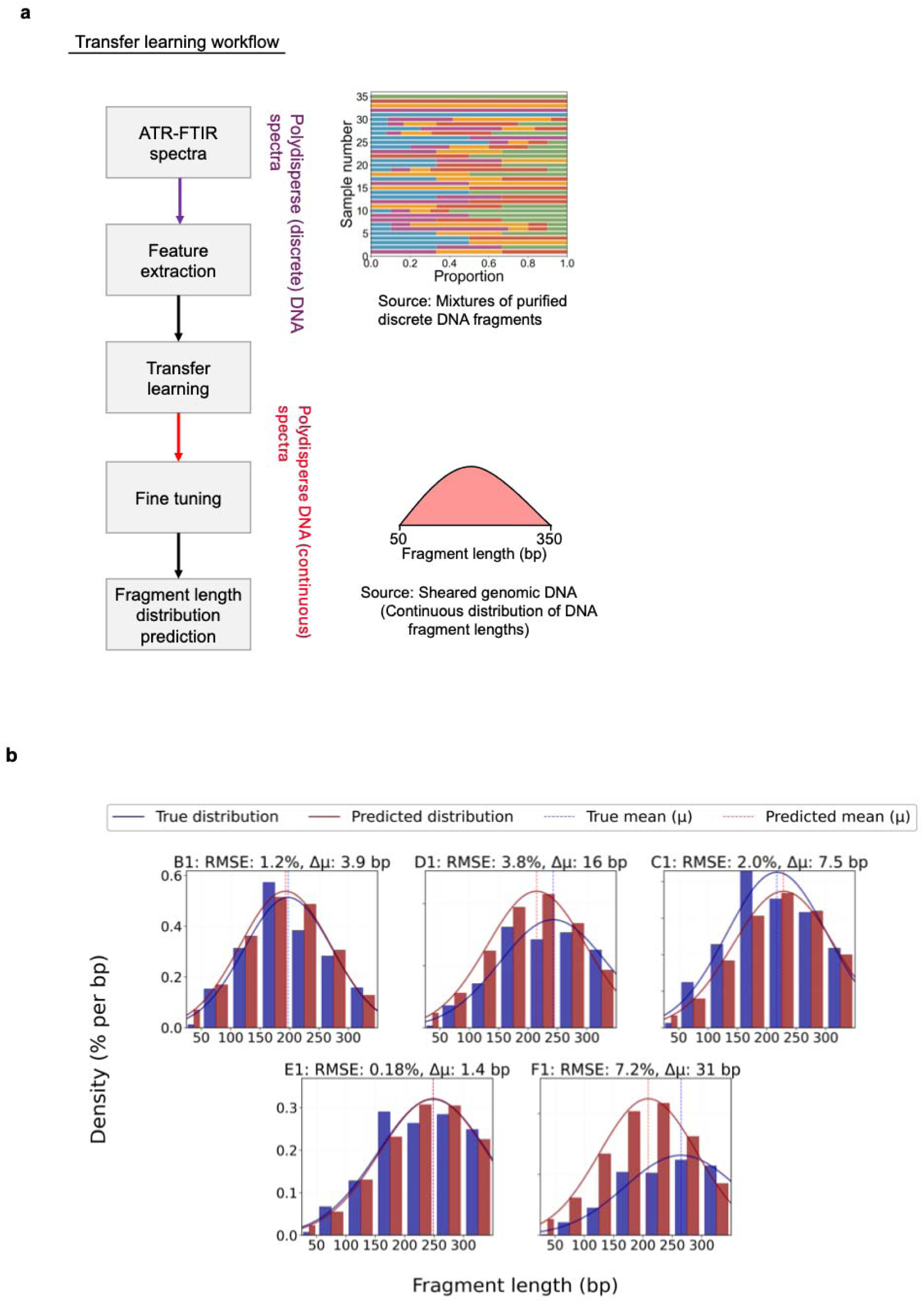
Predicting DNA fragment length in complex DNA polydisperse samples. **(a)** Transfer learning workflow for extending fragment length prediction from discrete polydisperse DNA mixtures to biological DNA samples containing continuous fragment length distributions. A 1D-CNN was pre-trained on pure DNA ATR-FTIR spectra with augmented spectra acquired from mixtures of purified DNA fragments with discrete lengths combined in defined proportions. The pre-trained network was then repurposed as a feature extractor, and model weights were fine-tuned using a small set of biological samples containing sheared genomic DNA, enabling adaptation from the purified domain (discrete fragment predictions) to the biological domain (continuous fragment length distributions) and prediction of fragment length distributions in heterogeneous DNA samples.**(b)** Testing on the independent test samples. The transfer learning approach yields good alignment between the predicted (red) and expected (blue) base pair distributions.

The transfer learning model was trained to perform multi-output regression, predicting the proportion of DNA across seven fragment length bins (25-50 bp, 50-100 bp, 100-150 bp, up to 300-350 bp) using the reference data obtained via gel electrophoresis for these same samples. The fine-tuned model was then applied to the averaged ATR-FTIR spectra from the five unseen biological samples (B1-F1) in the test set. The compositions of these samples are detailed in Supplementary Table 5. As shown in Figure 6b, the transfer learning framework delivered good predictive performance across all of the test samples. The model achieved overall error for the mixtures, with the highest RMSE 2.3% and lowest being 1.3%, and the average absolute shift in distribution centres between the predicted and true calculated value (Δμ) was approximately 7 bp.

In summary, these results show that transfer learning enables spectral models trained on spectra of purified DNA and derived augmented spectra, to predict fragment length profiles in biological samples containing polydisperse DNA. Achieving accurate predictions in complex biological samples, rather than just controlled mixtures, demonstrates the practical viability of this approach for fragmentomic profiling.

## Discussion

In this study we demonstrate for the first time that vibrational spectroscopy combined with machine learning can be used to quantify DNA fragment length distributions in complex samples without prior separation or labelling. We provide proof of principle that deep learning models can disentangle and learn fragment specific spectral features from purified DNA and transfer this knowledge to predict distributions in complex samples, achieving high accuracy.

DNA fragment length estimation is a key component of several molecular biology applications, including in next generation sequencing library preparation ^1,33,34^. DNA fragment length profiles in blood can also serve as biomarkers for monitoring therapy response and disease progression in cancer, as circulating tumour DNA molecules are typically shorter than those derived from normal cells. ^4,31,35^. Currently, DNA fragment length is assessed using gel electrophoresis-based systems, which rely on physical sample separation using migration speed. These technologies are expensive, require significant instrumentation, and are destructive to the sample being analysed. As a potential alternative, vibrational spectroscopy offers a rapid, non-destructive portable approach for DNA characterisation. Previous vibrational spectroscopy studies were focused on qualitative discrimination or structural characterization ^36–38^, with recent quantitative work predicting DNA base content from spectral changes ^39,40^.

Our approach presents an advancement in the field, where we introduce a methodology for DNA fragment length quantification using vibrational spectroscopy. For monodisperse fragment quantification, we developed machine learning models using ATR-FTIR and Raman spectroscopies on their own and then by integrating the complementary vibrational data from both spectroscopy techniques using data fusion. The data fusion model showed improved performance compared to individual spectroscopy model and quantified individual fragment lengths with an RMSE of 17 bp.

Each vibrational spectroscopy modality revealed length dependent differences in both spectral intensity and wavenumber shifts, primarily across the phosphate backbone, nucleobases and hydrogen bonding regions. In the ATR-FTIR spectra, intensity changes were most pronounced at ∼1080cm^-1^ with wavenumber shifts at ∼1250cm^-1^. In Raman spectra, length-dependent intensity differences were most significant at 1085cm^-1^ and 1450cm^-1^, with wavenumber shifts at ∼1250cm^-1^. Among these features, phosphate backbone associated bands emerged as robust indicators of fragment length. This is likely because phosphodiester bond content scales linearly with DNA fragment length, minimally influenced by sequence composition. In contrast, nucleobase associated regions are expected to be strongly influenced by sequence composition and therefore less suitable as a standalone fragment length feature.

Building on these findings, we extended our approach to enable the simultaneous quantification of multiple DNA fragment lengths in polydisperse mixtures. For this more complex task, we focused exclusively on ATR-FTIR spectroscopy as fragment length dependent spectral changes were more clearly resolved compared to Raman, allowing more accurate prediction of mixture compositions. In addition, ATR-FTIR offers practical advantages for clinical translation, as a single modality reduces instrumentation complexity and costs.

A key innovation of this study lies in the application of transfer learning to bridge the gap between purified DNA and biological samples. A 1D-CNN trained on augmented spectra derived from 35 purified DNA mixtures learned spectral features associated with fragment length and accurately predicted fragment proportions with an average RMSE of 6.5%. Fine-tuning this pre-trained model on a cohort of six DNA samples with a continuous length profile, allowed it to adapt to spectral differences of more complex biological samples. Importantly, the method requires only 4 μl of sample and 15 minutes of passive drying, with no consumables beyond cleaning materials, and allows full sample recovery for downstream applications.

These results should be interpreted within the study’s constraints. First, the models do not incorporate sequence specific information, hence differences in base composition between fragments of the same length may influence predictions. Second, the purified training set was limited to 50-300 bp, restricting analysis to the 300-350 bp region despite reference (gold standard) detection up to ∼1500 bp. Third, the biological cohort was small (n=11), with six samples used for transfer learning and five for blind testing. While this was sufficient to demonstrate feasibility across distributions, this cohort is limited relative to expected variability in DNA samples.

In conclusion, this study provides the first demonstration that vibrational spectroscopy enables quantitative DNA fragment length analysis in solution, establishing a rapid, non-destructive analytical platform. Importantly this capability could form a foundation for scalable fragment length assessment in genomic workflows and circulating tumour DNA analysis.

## Methods

### DNA Samples

Purified DNA fragments of defined lengths 50 bp (catalogue number SM1421), 100 bp (SM1441), 150 bp (SM1601), 200 bp (SM1611), and 300 bp (SM1621); 0.5μg/μl (in TE buffer) were purchased from ThermoFisher Scientific. For the monodisperse experiments, these samples were used in their undiluted form and were directly pipetted to the ATR crystal for spectral acquisition. To generate a comprehensive dataset for 1D-CNN model development, polydisperse DNA mixtures were prepared from the purified DNA fragments using an extreme vertices experimental design. This design strategy, implemented through Minitab software, explored the five-fragment mixture space by efficiently sampling combinations that would enable accurate quantification of proportions. The resulting 35 sample design set comprised: (i) pure component samples, (ii) binary mixtures containing two fragments in equal or unequal proportions, (iii) ternary mixtures with three fragments, (iv) quaternary mixtures with four fragments, (v) centroid mixtures with all five fragments in equal proportions, and (vi) additional mixtures with all five fragments in varying proportions. The coverage from one component samples to multi fragment mixtures allowed comprehensive model training and evaluation across diverse fragment distribution profiles. Each mixture was prepared by the volumetric combination of fragment stock solutions at a fixed total volume of 12 μl. Since all stock solutions were obtained at identical concentrations, the relative fragment abundance in each mixture was defined by the volumetric ratios used during mixing, with exact compositions reported in Supplementary Table 3.

Rat genomic DNA was prepared using the Monarch® HMW DNA extraction kit for tissue (NEB #T3060) following manufacturer’s instructions. Rat mammary tissues were pestle-homogenised and lysed in HMW gDNA tissue lysis buffer and proteinase K (600 μl lysis and 20 μl proteinase K per 15mg tissues) at 56°C for 5 h agitated at 1400 rpm in a thermo mixer. The lysate was then digested with 10 μl of RNase A at 56°C for 10 min agitated at 1400rpm. 300 μl of protein separation solution was added and the tube was inverted for 1 min and spun for 10 min at 16000g. The upper phase was transferred using a 1000 μl wide-pore pipette tip to a new 2 ml tube and 2 DNA capture beads and 550 μl isopropanol were added. The tube was slowly inverted for 30 times allowing 5-6 seconds for each inversion. The liquid was discarded and 500μl of gDNA wash buffer was added to the beads. The tube was inverted for 3 times and the wash buffer was removed. This wash step was repeated once. The beads were poured onto a bead retainer in a collection tube. After the tube was spun briefly to remove residual wash buffer. The beads were poured from the retainer into a new 2 ml tube. 100 μl elution buffer II was added to the beads and incubated for 5 min at 56°C agitated at 300 rpm in a thermal mixer. The beads and elution volume were poured onto the saved retainer in a 1.5 ml tube. The tubes were spun at 12000g for 30 sec. The eluate was saved and resuspended up and down with a wide-pore tip for 10 times. This genomic DNA was subsequently sheared using Covaris S220 following manufacturer’s instructions. Briefly, 5μg of DNA in 0.1M Tris-HCl (pH8.0) was put in a microTUBE-50 (Covaris PN520166) in a final volume of 55 μl for each condition. The water level was set at 10 at 7°C. Target BP (Peak) was set to 150, 200, 250, 300 or 350 bp. The fragment length distribution for each condition was analysed using the Agilent 4200 TapeStation system with HS D1000 ScreenTape.

### ATR-FTIR spectroscopy

ATR-FTIR spectra were acquired using an Agilent Cary 670 FTIR spectrometer equipped with a liquid nitrogen cooled MCT detector and a MIRacle 9 bounce reflection ATR fitted with a diamond coated ZnSe crystal. For each measurement, 4 µl of DNA was pipetted onto the ATR crystal and allowed to air dry at room temperature for approximately 15 min to form a uniform thin film. Spectra were collected across the 700-6000 cm^-1^ range at a spectral resolution of 4 cm^-1^, with 64 scans averaged per acquisition to enhance the signal to noise ratio. Each sample was measured in triplicate, with the ATR cleaned between each measurement and nine repeated scans performed per sample before proceeding to the next composition.

### Raman spectroscopy

Raman measurements were performed using a Renishaw inVia micro-Raman spectrometer equipped with a 532 nm excitation laser and a 20× objective lens. For sample preparation, 2 μl of each DNA fragment length solution was deposited onto a CaF substrate and dried at room temperature. CaF was chosen due to its low fluorescence background in the biologically relevant spectral region, thereby improving spectral quality.

Spectra were collected over the 600-1800 cm^-1^ range. After focusing the microscope objective on the dried deposit, Raman spectra were acquired at the outer edge of the sample using 10 accumulations with a 10 s integration time, yielding 15 spectra per position. The laser position was then adjusted, and an additional 15 spectra were collected to ensure adequate coverage of the dried sample’s surface.

### Data preprocessing

All preprocessing procedures were carried out after splitting the data into training and testing subsets to prevent data leakage. Optimised parameters derived exclusively from the training data were subsequently applied to the test set. For ATR-FTIR spectra, the analysis was truncated to 750-1800 cm^-1^, as the diamond ATR crystal introduces strong absorbance between ∼1800-2700 cm^-1^, limiting interpretability in this region. For Raman spectra, the full acquired wavenumber range was retained without truncation and cosmic ray artifacts were removed using Renishaw’s WiRE software. Vector normalization was then performed across the most informative spectral regions (FTIR: 750-1800 cm^-1^ and Raman: 600-1800 cm^-1^) to account for intensity variations. A manual three-point baseline correction was applied to all spectra to eliminate background drift, followed by a Savitzky-Golay second order derivative transformation to enhance spectral resolution and resolve overlapping features.

### Machine learning

PLSR was used as a supervised learning approach to quantify individual DNA fragment lengths from their corresponding spectra. This method is particularly well suited for high dimensional spectroscopic data, as it effectively addresses multicollinearity by extracting latent variables that maximise the covariance between the features and the target response variable (fragment length).

To prevent data leakage, data splitting was performed in a manner that respected the structure of our measurements. In spectroscopic experiments, replicates refer to repeated measurements of the same kind of sample following physical replacement or repositioning (e.g., removing a DNA sample from the ATR crystal and reapplying it). In contrast, repeats refer to multiple spectra acquired from the same physical sample without repositioning (e.g., multiple spectra collected from the same DNA spot). Since repeats from the same replicate originate from the same underlying sample, they are not statistically independent and were therefore kept together during train/test splitting to avoid overestimating model performance.

For the FTIR dataset, data splitting was performed by on a replicate basis where the first two replicate sets were used for training and cross-validation and the third was held out as an independent test set. For the Raman dataset, splitting was based on physical sampling location to prevent data leakage. For each sample measured, spectra acquired around one laser spot were used exclusively for model training, whereas those from the proximity of a second spot were reserved for independent testing.

Model optimisation and latent variable selection were performed on the training set to prevent overfitting with the optimal number of PLS components being determined using 10-fold cross validation in combination with Van der Voet’s F test. This method objectively compares the cross-validation prediction errors (mean squared error) between models with *n* and *n + 1* components. The first non-significant comparison (*p* ≥ *0.05*) was taken as the criterion for selecting the optimal number of components, ensuring a parsimonious model that captures relevant variance without fitting to noise.

The final regression model performance was assessed using the coefficient of determination (R^2^), reflecting the proportion of variance explained by the model, and the root mean square error (RMSE), which evaluates the prediction error.

### Convolutional Neural Network (CNN) development

All deep learning model development, including baseline training, hyperparameter optimization, and transfer learning, was conducted on the University of Southampton’s high-performance computing (HPC) cluster, Iridis X. Training jobs were executed on NVIDIA L4 GPU nodes using the CUDA-enabled TensorFlow backend, with cuDNN providing GPU-accelerated operations. Reproducibility was ensured using deterministic TensorFlow settings, fixed random seeds, and strict group-wise data splits that kept all replicates from each sample together, preventing data leakage and ensuring test performance reflected true generalization to unseen samples.

Hyperparameter optimization was performed using the Hyperband algorithm to efficiently explore learning rates, convolutional filters, kernel sizes, dropout levels, regularization strengths, dense layer architectures, neuron counts, and data augmentation parameters. This approach allocated computational resources to the most promising configurations that minimized validation loss.

To increase model robustness, the purified DNA training set was augmented six-fold using controlled perturbations (Supplementary Fig. 7) inspired by the spectral variation framework of Bjerrum et al. (2017) ^41^. Realistic instrumentation variation was simulated through three transformation types applied in combination: (i) intensity scaling by multiplying each spectrum by a uniform factor of 0.99-1.01 times the training set standard deviation, (ii) vertical offset implemented as an additive shift sampled between - 0.01 and +0.01 absorbance units scaled by the spectral training set standard deviation, and (iii) uniform noise introduced as random perturbations up to 0.001 absorbance units, scaled by the training set standard deviation. To enforce physically meaningful outputs, all model predictions were passed through a custom *Constrained Prediction Layer* that normalized outputs to sum to 100%.

The optimized architecture consisted of convolutional layers, dense hidden layers, and a linear output layer predicting five fragment percentages. The model was trained using the Adam optimizer with Huber loss (Supplementary Fig. 8) and a 20% validation split. To enforce physically meaningful outputs, all model predictions were passed through a custom *Constrained Prediction Layer* that normalized outputs to sum to 100%.

### Transfer learning for biological samples

A two-stage transfer learning framework was developed to adapt spectral representations learned from polydisperse (discrete) DNA mixtures to polydisperse (continuous) biological DNA samples. In Stage 1, the pretrained 1D-CNN served as a feature extractor with all convolutional layers (including weights, biases, and regularization parameters) frozen to preserve learned DNA features. A new prediction head comprising dense layers was trained on top of the frozen feature extractor, allowing the model to learn biologically relevant mappings from purified fragment features to real DNA fragment proportions without disrupting the pretrained convolutional filters.

After Stage 1 convergence, all layers were unfrozen in Stage 2 and the entire network was fine-tuned end-to-end using a reduced learning rate (0.0001 vs. 0.001). This fine-tuning allowed convolutional filters to adapt to biological sample complexity while retaining the learned structural features, thereby improving generalization.

Model optimization employed a customized Wasserstein-MSE loss function combining the Wasserstein distance and mean squared error (Supplementary Fig. 11). The Wasserstein component (Earth Mover’s Distance) considered the spatial ordering of fragment length intervals, penalizing predictions that misplaced mass across distant bins, while the MSE component penalized bin-specific errors, together encouraging accurate recovery of both interval-wise magnitudes and overall distributional shape.

### Feature selection and interpretation

To identify the most informative spectral regions contributing to DNA fragment length quantification, we examined the loading plots from the PLS-DA models. Loadings were plotted against wavenumber to visualise the magnitude and direction of influence for each spectral variable. High absolute loading values indicated wavenumbers that strongly drove the separation between fragment length classes in the latent space. This analysis was used to validate the optimal spectral window (750-1800 cm^-1^) for subsequent machine learning models, ensuring that the analysis focused on meaningful biological signals (phosphate, sugar, and nucleobase vibrations) while excluding regions dominated by noise and buffer contributions.

### DNA fragment length distributions

Fragment length distributions for the biological DNA samples were generated through shearing using a Covaris S200 instrument. Instrument parameter sets (TP settings) were used to target specific modal fragment lengths, including TP150, TP200, TP250, TP300, and TP350. For example, the TP150 setting was configured to produce a dominant fragment population centred around approximately 150 bp, while progressively higher TP settings were used to shift the fragment length distributions towards larger sizes.

Gel electrophoresis was used to check the resulting fragment length distributions and served as the reference measurement. Fragmentation experiments were repeated once and produced comparable electropherograms across duplicate preparations for all TP settings, with the exception of TP350, which exhibited broader distributions and increased inter-replicate variability.

### Prediction and visualization of fragment length distributions

For the purified DNA mixtures, the 1D-CNN was trained to predict the percentage contribution of each discrete fragment length (50, 100, 150, 200, 300 bp). For the biological data, the transfer learning model instead predicted the percentage of DNA assigned to predefined length intervals (25-50 bp to 300-350 bp) from the gel electrophoresis data. In both cases, the raw model outputs are therefore binned proportion vectors. To obtain a continuous representation of the underlying fragment length distribution, we converted these discrete proportions into a unimodal density estimate by treating them as weighted samples in length space, computing the weighted mean and standard deviation, and fitting a single Gaussian distribution. This fitted Gaussian was then evaluated on a fine grid of fragment lengths to generate a smooth curve for plotting, while the original discrete proportions were retained for calculating metrics such as RMSE, R^2^ and other bin wise performance metrics. Thus, both the purified (discrete fragment) and biological (interval based) predictions were handled in a unified framework that operates on discrete outputs but visualises them as continuous distributions.

### Data integration (fusion)

For each sample, we extracted the informative window from both modalities (IR: absorbance 750-1800 cm^-1^ and Raman: counts 600-1800 cm^-1^). Each dataset was independently normalized (min-max normalisation within its own wavenumber range) to prevent one modality from dominating due to scale differences. We then performed true axis concatenation where the IR wavenumber axis was kept in its original wavenumber range (750-1800 cm^-1^), and the Raman axis was shifted by a constant offset so that it begins immediately after the IR band, creating a single combined axis with no overlapping. Specifically, the shift is given as concat_shift = (max IR wavenumber - min Raman shift). The fused CSV file thus contains a continuous spectral vector in which the IR section is followed immediately by the shifted Raman section. To preserve physical interpretability, we also stored the original (unshifted) wavenumber/shift in an OrigWavenumber column and tagged each row with Source ∈ {IR, Raman}.

## Acknowledgements

This work was supported by the UK Engineering and Physical Sciences Research Council (EPSRC grant EP/S03109X/1). R.F thanks EPSRC DTP PhD studentship. We thank Breast Cancer Now for funding this work as part of Programme Funding to the Breast Cancer Now Toby Robins Research Centre. S-J.S. is a Lister Institute Prize Fellow.

## Author information

G.S.M. and S-J.S. conceptualized and directed the research. R.F. designed and performed the DNA fragment length experiments, developed and implemented the machine learning models, and performed the primary data analysis. G.S.M. provided funding and overall project supervision. W.A. contributed to experimental design and interpretation of results. I.S. prepared and characterised the biological samples. All authors contributed to manuscript writing and approved the final version.

## Data Availability

The data for this work are accessible through the University of Southampton Institutional Research Repository via https://doi.org/10.5258/SOTON/D3838 (2026).

## Ethics declarations

### Competing interests

The authors G.S.M, S-J.S and R.F are inventors of a patent application related to methods for analysing DNA samples (UK priority patent application GB2414133.5, filed 26 September 2024).

### Additional Information

Supplementary Information is available for this paper.

