## Supplementary Information for "DNA fragment length analysis using machine learning assisted vibrational spectroscopy"

**Supplementary Table Legends:**

**Supplementary Table 1:** ATR-FTIR peak assignments for spectra. Assignments for the identified peaks in the ATR-FTIR spectra shown in Figure 2a of the main text.

**Supplementary Table 2**: Raman peak assignments for spectra. Assignments for the identified peaks in the Raman spectra shown in Figure 2b of the main.

**Supplementary Table 3**: Composition of purified DNA fragment mixtures used for model development. DNA fragments of defined lengths (50, 100, 150, 200 and 300 bp) were combined using an extreme vertices experimental design to generate 35 single, binary, ternary, quaternary and five component mixtures. All mixtures were prepared by volumetric combination of equal concentration stock solutions (0.5 μg/μl in TE buffer) to a fixed total volume of 12 μl. Exact volumetric ratios and corresponding fragment compositions for each mixture are reported in the table. The weighted mean fragment length ($\bar{L}$) for each purified mixture was calculated as:

$\bar{L}=\frac{\sum_{i=1}^{N} v_{i}m_{i}}{\sum_{i=1}^{N} v_{i}}$	(1)

where $v_{i}$ is the volume (μl) of fragment size i, and mᵢ is the midpoint fragment length (bp) for that size. This equation weights each fragment size by its volumetric proportion in the mixture.

**Supplementary Table 4**: Continuous fragment length distribution profiles for biological DNA samples used for model training. Fragment percentage (%) across defined fragment length bins (25-350 bp) derived from gel electrophoresis electropherograms for biological DNA samples containing continuous fragment length distributions. Values represent bin-wise fragment distributions for the training cohort samples (B1-G1). The calculated weighted mean ($\bar{L}$) fragment lengths for the training cohort were found using the following equation:

$\bar{L}=\frac{\sum_{i=1}^{N} p_{i}m_{i}}{\sum_{i=1}^{N} p_{i}}$	(2)

Where *pᵢ* is the percentage of fragments in bin *i*, and *mᵢ* is the midpoint of that size range. The denominator normalizes for incomplete distribution coverage (fragments outside the measured 25-350 bp range). This resulted in the following weighted means 178 bp (B1), 208 bp (C1), 205 bp (D1), 216 bp (E1), 226 bp (F1), and 229 bp (G1). Additionally, the median was calculated as:

$Median = L + \left( \frac{\left( 0.5 \sum_{i=1}^{n} p_{i}- C_{prev} \right)}{p_{bin}} \right)w$	(3)

where *L* is the lower bound of the fragment size bin containing the median, w is the bin width, $p_{bin}$ is the percentage within the median bin, $C_{prev}$ is the sum of all the percentage up to the median bin, and $\sum_{i=1}^{n} p_{i}$ denotes the total percentage across all bins for a given sample. This equation accounts for cases where the binned fragment-size distributions do not sum to 100%, therefore the median fragment length is conditional to the analysed size range. Using equation (3) resulted in the following found median values (B1) 174 bp, (C1) 204 bp, (D1) 200 bp, (E1) 216 bp, (F1) 230 bp, (G1) 234 bp.

**Supplementary Table 5**: Quantitative fragment length distribution profiles for biological DNA samples used for independent model testing. Fragment percentage (%) across defined fragment length bins (25-350 bp) derived from gel electrophoresis electropherograms for biological DNA samples containing continuous fragment length distributions. Values represent bin-wise fragment distributions for the independent test cohort samples (B1-F1). The calculated weighted mean fragment lengths for the independent test cohort, using equation (2) were 196 bp (B1), 209 bp (C1), 224 bp (D1), 226 bp (E1), and 234 bp (F1). The medians for this test set were (B1) 190 bp, (C1) 206 bp, (D1) 226 bp, (E1) 229 bp, (F1) 242 bp.

**Supplementary Figures**

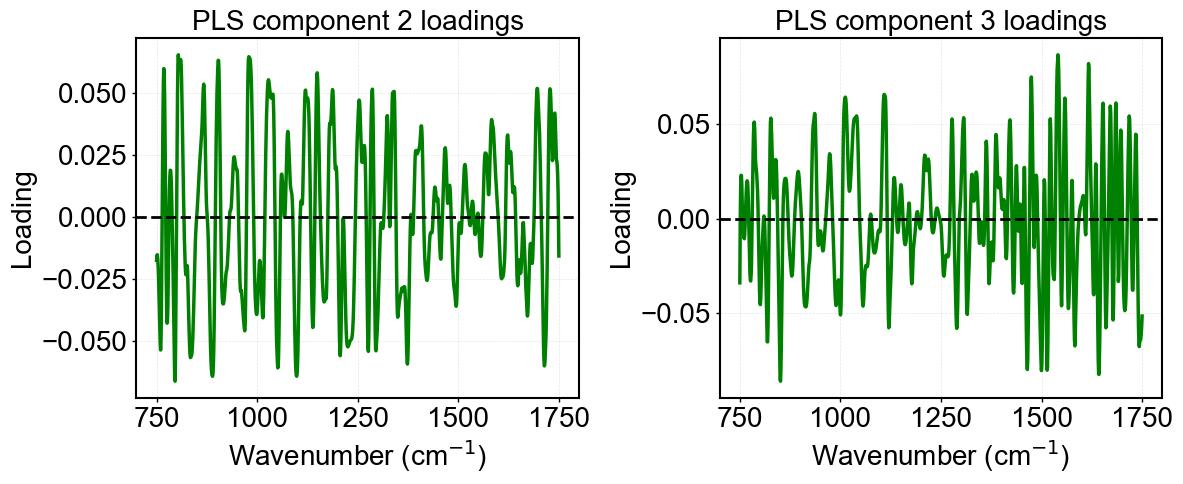

**Supplementary Figure 1**: ATR-FTIR spectra PLS-DA loading plots. Loading vectors for PLS components 2 (LV2) and PLS components 3 (LV3) from the ATR-FTIR spectra for the different individual DNA fragment lengths. The plot shows the loadings across the analysed wavenumber regions, indicating that LV2 captures better loadings between 800-1300cm^-1^ whereas for LV3 loadings this was from 1500-1800cm^-1^.

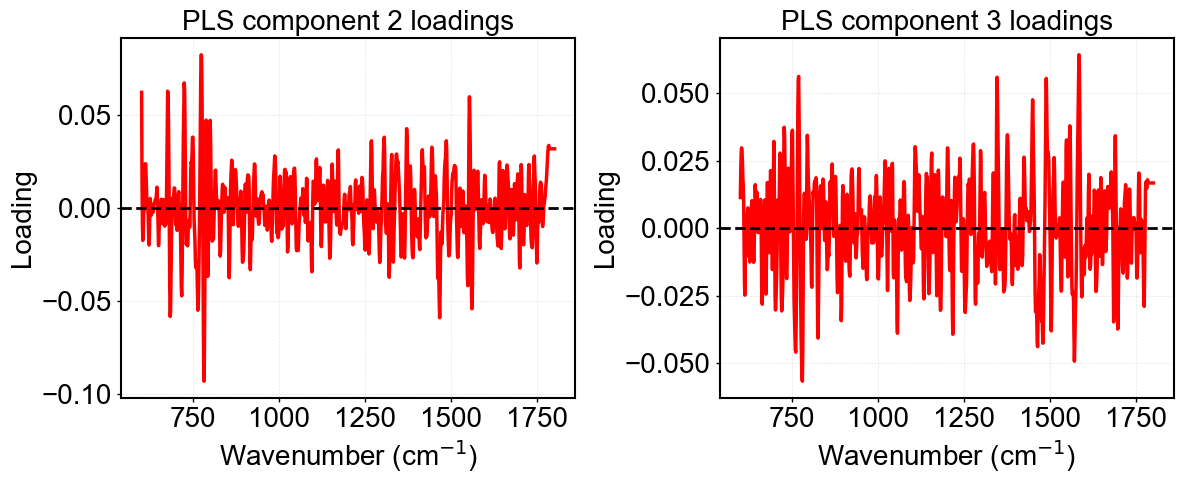

**Supplementary Figure 2**: Raman spectra PLS-DA loading plots. Loading vectors for PLS components 2 (LV2) and PLS components 3 (LV3) from the Raman spectra for different individual DNA fragment lengths. Highest loading values in LV2 were observed in the lower wavenumber 600-800cm^-1^ with the strongest at 785cm^-1^. Overall LV3 was nosier, but strong loadings were found occurring from 1300-1580cm^-1^.

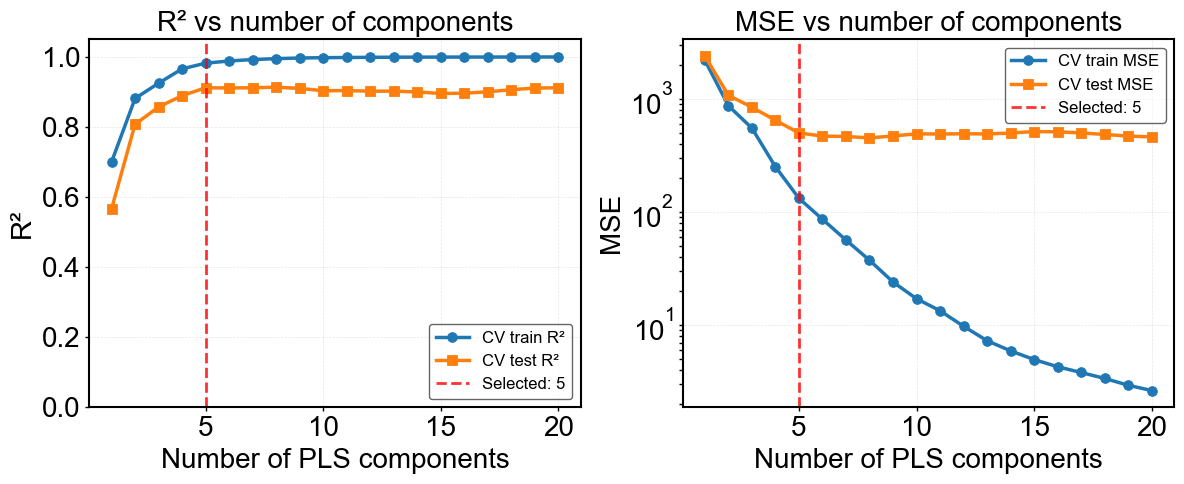

**Supplementary Figure 3**: Optimisation of the number of latent variables (LVs) for ATR-FTIR PLSR models of monodisperse DNA samples, evaluated using the coefficient of determination (R^2^) and mean squared error (MSE). The optimal LV number was determined by cross-validation on the CV train/test split, and model significance was assessed using Van der Voet’s F-test.

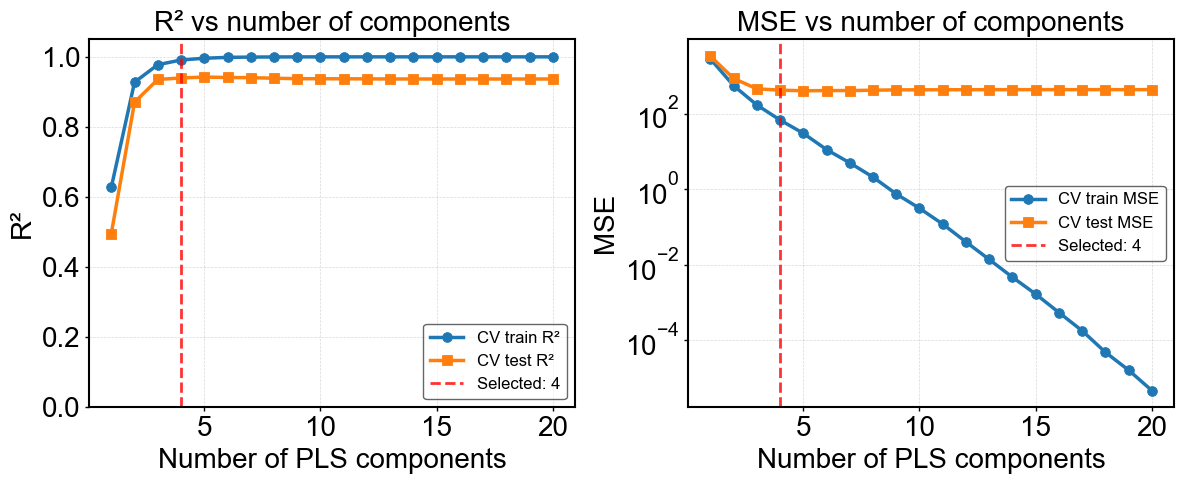

**Supplementary Figure 4**: Optimisation of the number of latent variables (LVs) for Raman PLSR models of monodisperse DNA samples, evaluated using the coefficient of determination (R^2^) and mean squared error (MSE). The optimal LV number was determined by cross-validation on the CV train/test split, and model significance was assessed using Van der Voet’s F-test.

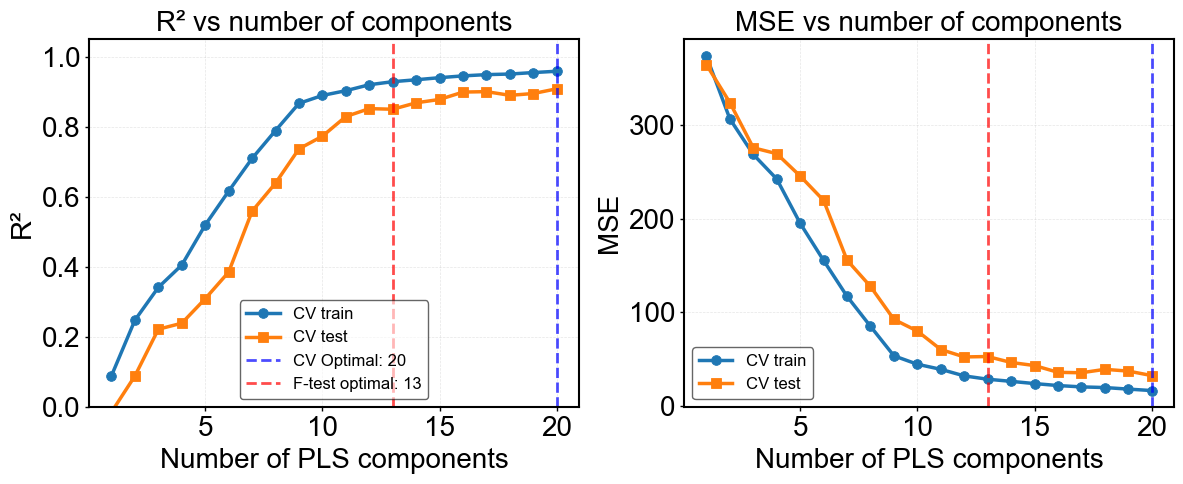

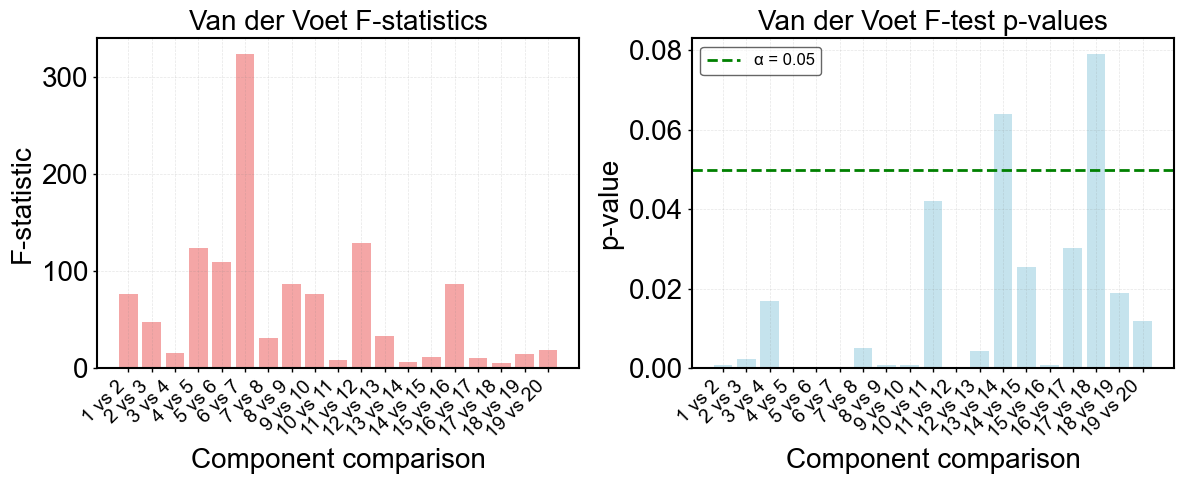

**Supplementary Figure 5**: PLSR DNA mixture latent variable optimization. LV optimisation for the PLSR model using R^2^ and MSE from the cross-validation. Here, CV train represents performance across the training folds, while CV test reflects performance on the validation folds. An F-test was applied to determine the optimal number of LVs. However, the requirement for a high number of LVs to explain model variance may indicate that the model is capturing noise rather than features truly associated with DNA fragment length. Hence, a 1D-CNN architecture was selected as a more suitable modelling approach.

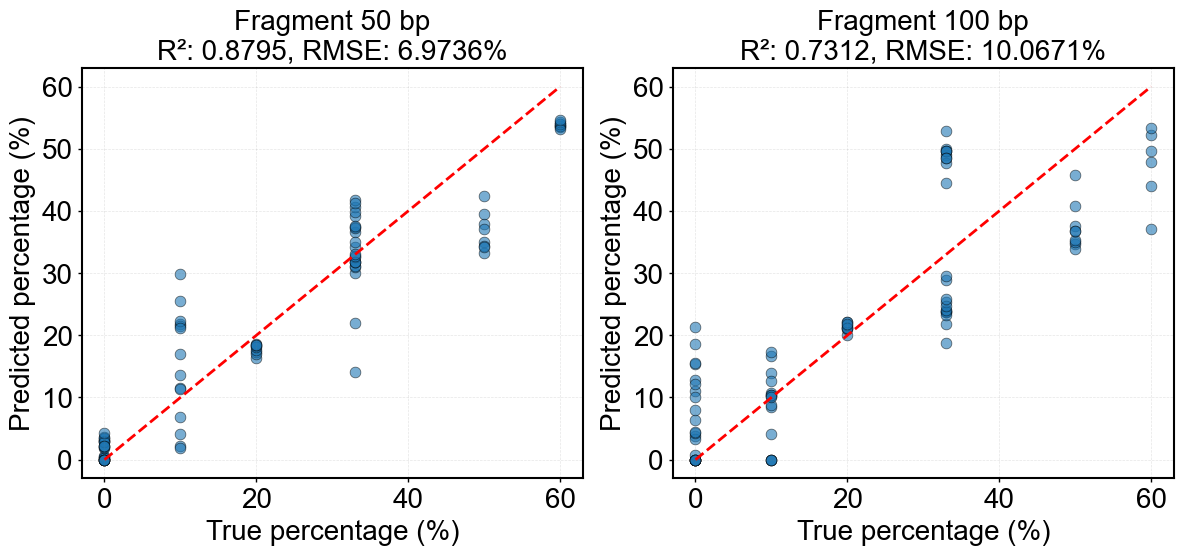

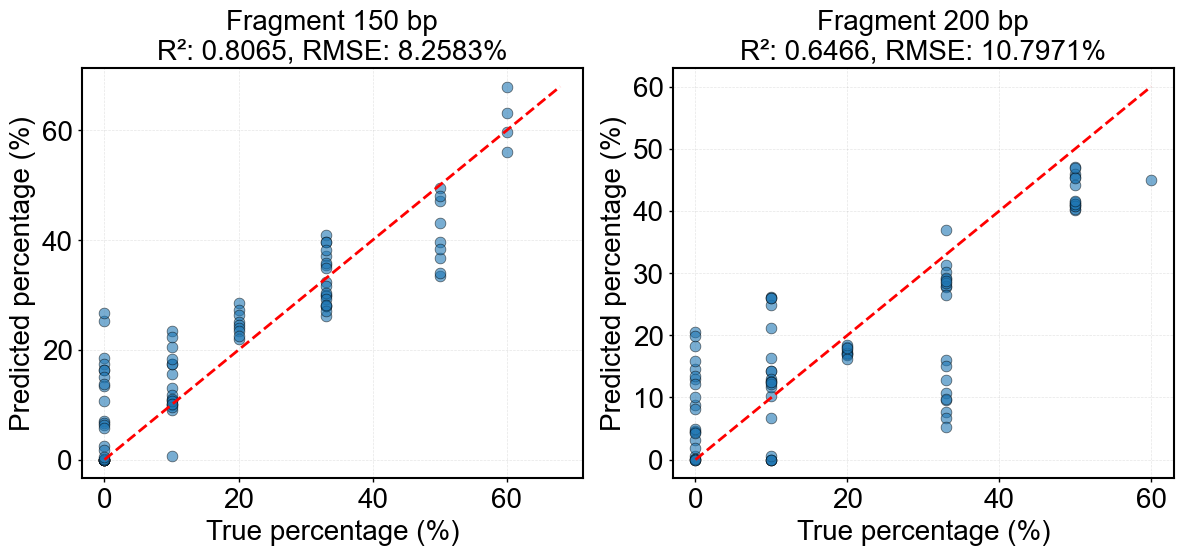

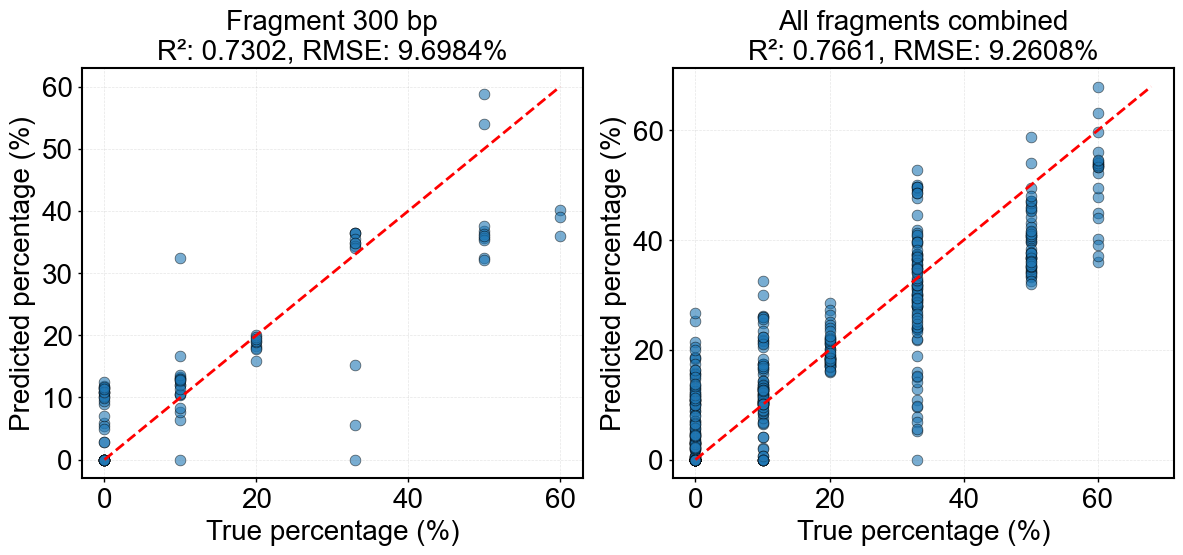

**Supplementary Figure 6**: PLSR predictions of fragment proportions in polydisperse DNA mixtures. The figure shows the performance of the PLSR model on pure DNA mixtures, summarising predicted versus true fragment-wise percentages across all mixtures for each DNA fragment length. PLSR was used as a baseline regression model. While good performance was achieved for shorter fragments, with the strongest results observed for the 50 bp fragment (R^2^ = 0.88, RMSE = 7.0 %), performance was worse for longer fragments, with the weakest performance for the 200 bp fragment (R^2^ = 0.65, RMSE = 9.7%). Overall performance across all fragments and mixtures produced an R^2^ of 0.76 and an RMSE of 9.3 %, motivating the use of a 1D-CNN model.

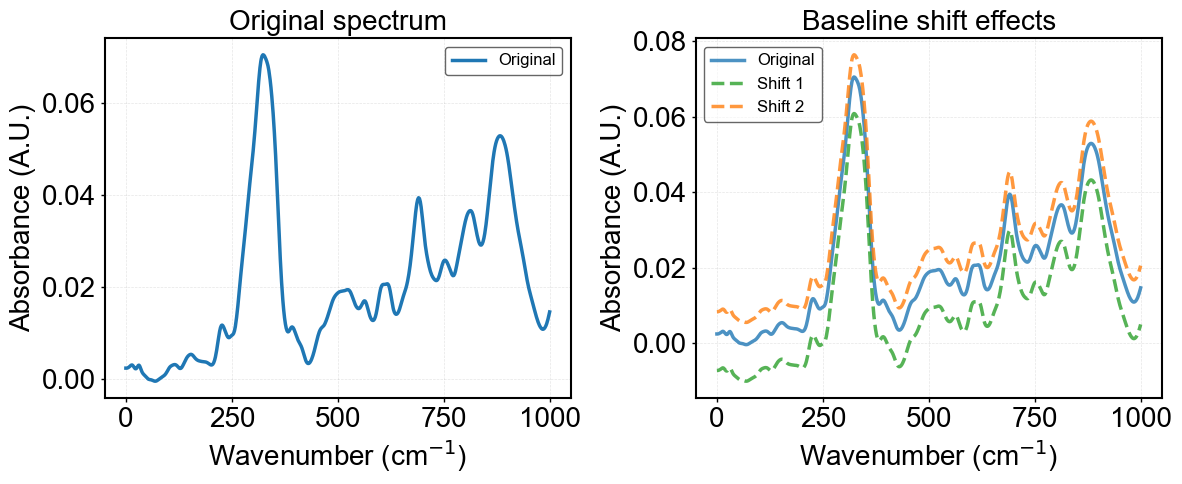

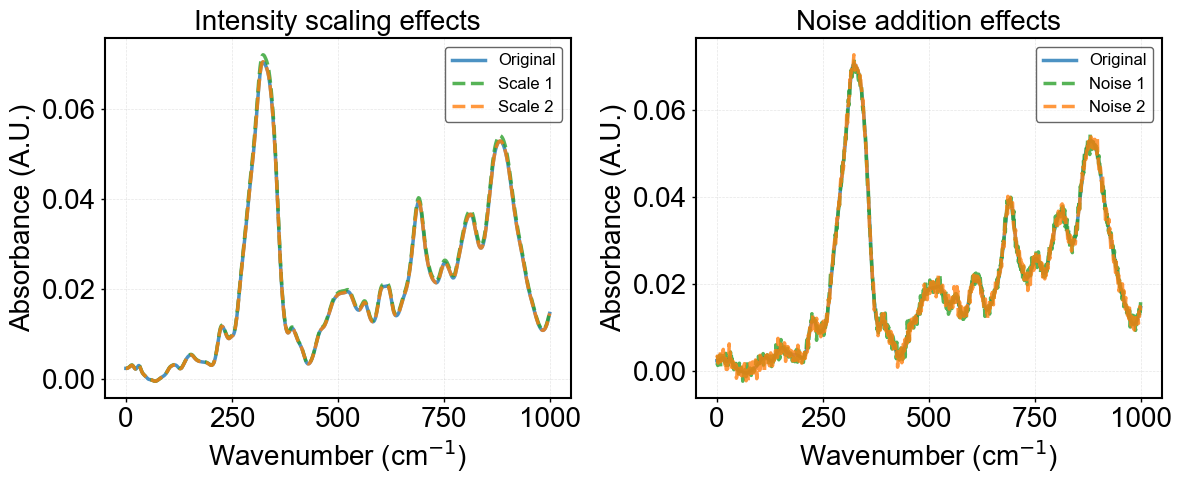

**Supplementary Figure 7**: Examples of data augmentation applied to ATR-FTIR spectra for the 1D-CNN model. The figure illustrates the different augmentation strategies applied to the original spectra. This included baseline shift augmentation, where spectra are vertically shifted to simulate baseline drift, intensity scaling augmentation, in which the overall spectral amplitude is scaled to mimic concentration or pathlength variations and noise addition augmentation, producing spectra with increasing levels of random noise to reflect instrument and environmental variability.

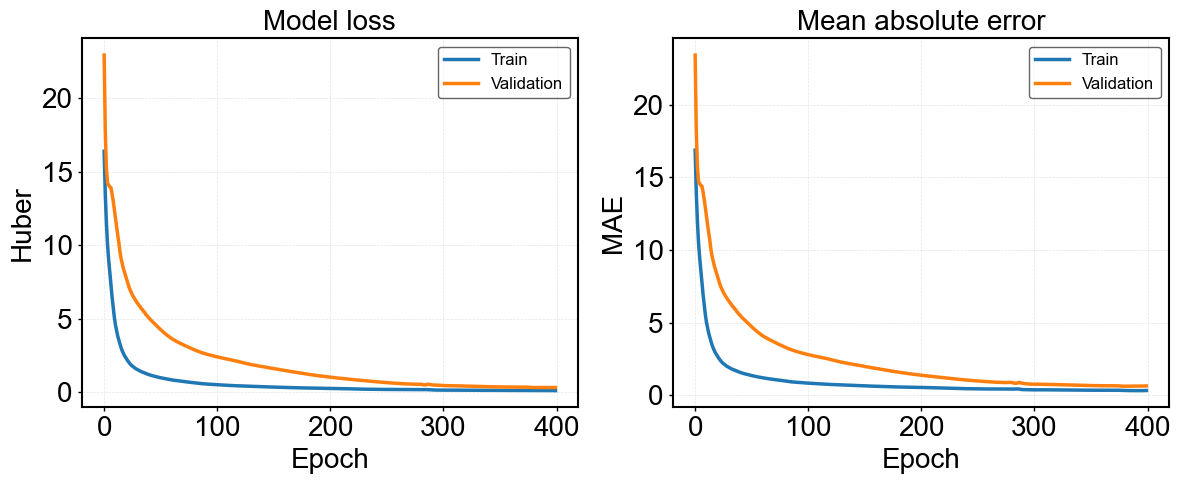

**Supplementary Figure 8**: 1D-CNN model training plot for polydisperse DNA mixtures. Figure shows the training history of the 1D-CNN model, monitored using both the Huber loss used for optimisation and the mean absolute error (MAE). In both metrics, the training and validation show a smooth and gradual convergence, reaching stability at approximately 350-400 epochs. After convergence, the validation curves remain consistently close to the training curves, with no fluctuations, indicating the absence of overfitting. The alignment of training and validation curves show stable and healthy learning.

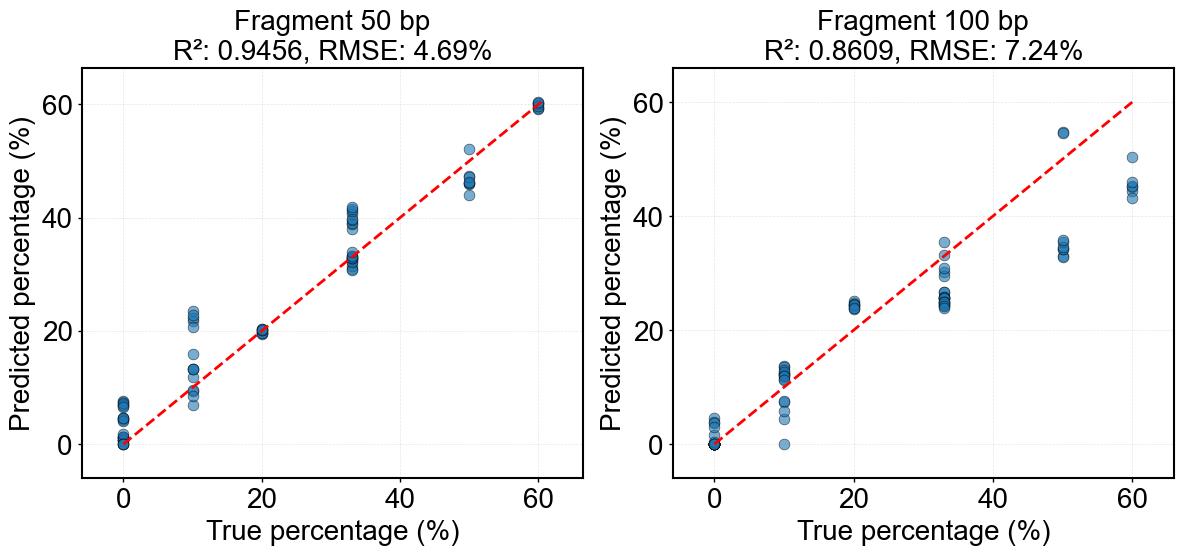

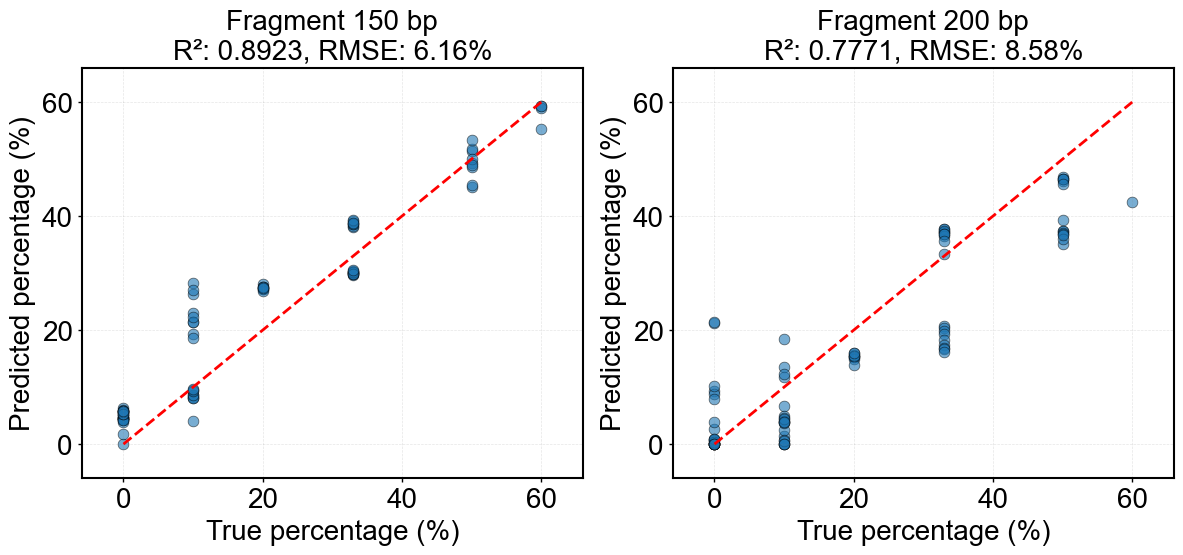

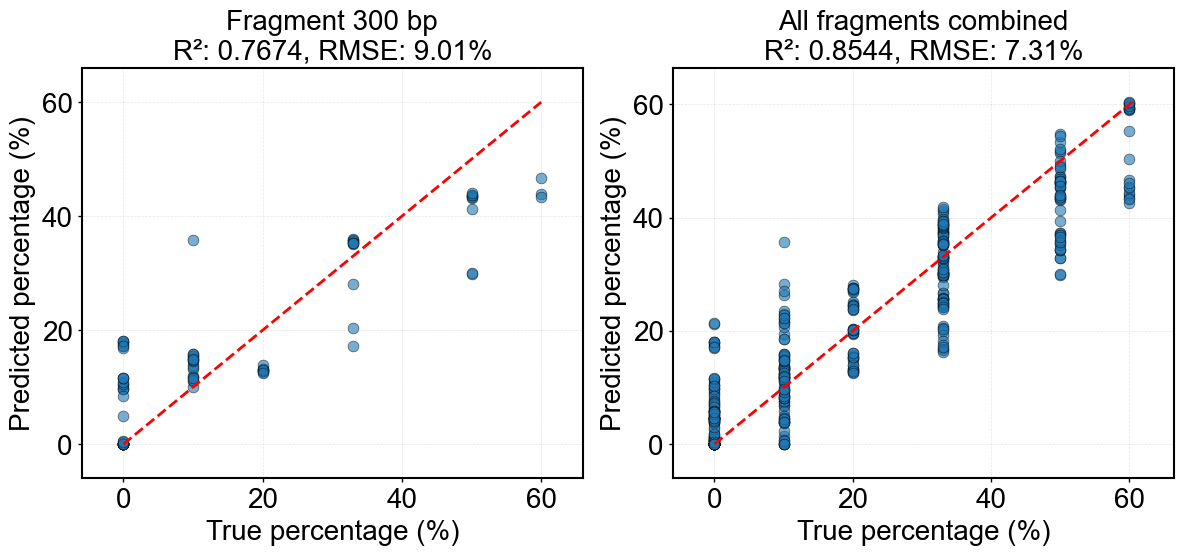

**Supplementary Figure 9**: 1D-CNN regression of DNA fragment percentages in synthetic DNA mixtures. The figure shows the performance of the 1D-CNN model in predicting fragment-wise DNA length percentages across the synthetic mixtures. The strongest performance was observed for the 50 bp fragment (R^2^ = 0.94, RMSE = 4.7%), while the weakest performance occurred for the 300 bp fragment (R^2^ = 0.77, RMSE = 9.0 %). Across all fragments and mixtures combined, the model achieved an overall R^2^ of 0.85 and an RMSE of 7.3 %, representing a moderate improvement over the PLSR baseline.

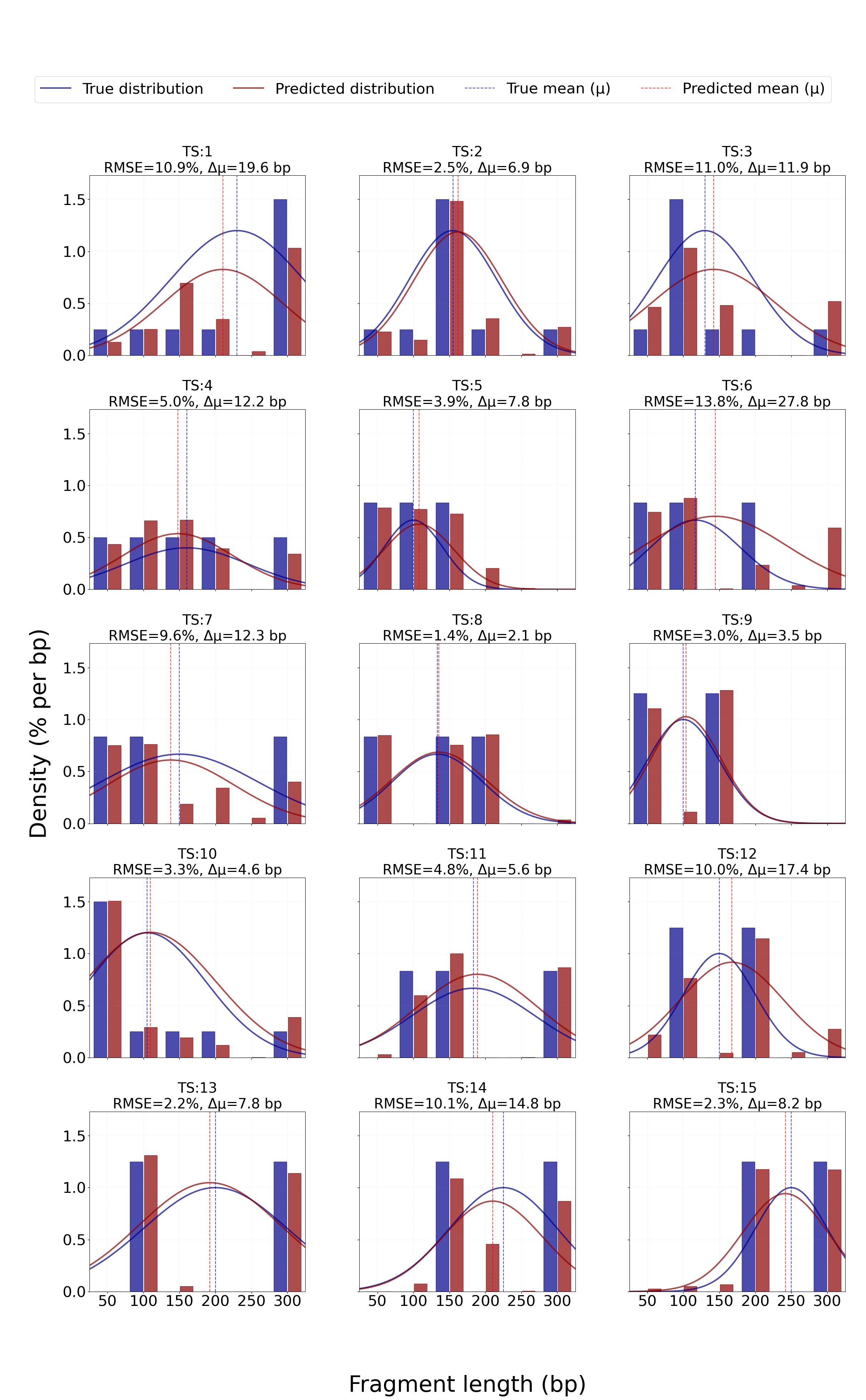

**Supplementary Figure 10**: 1D-CNN predictions of fragment length distributions for held-out test samples. The figure presents the complete set of polydisperse DNA mixtures held out for independent evaluation using the 1D-CNN model. Each plot shows the true fragment length distribution (blue) and the corresponding model-predicted distribution (red), overlaid with Gaussian fits and vertical lines indicating the weighted mean fragment lengths (μ). Model performance for each mixture is summarised using the distribution RMSE (%) and the weighted mean error (Δμ, bp). All mixtures shown were fully independent of model training and optimisation. Across the held-out test set, the model demonstrates consistent reconstruction of overall fragment length distributions, with most mixtures exhibiting low RMSE and small weighted mean deviations, indicating good generalisation to unseen polydisperse DNA compositions.

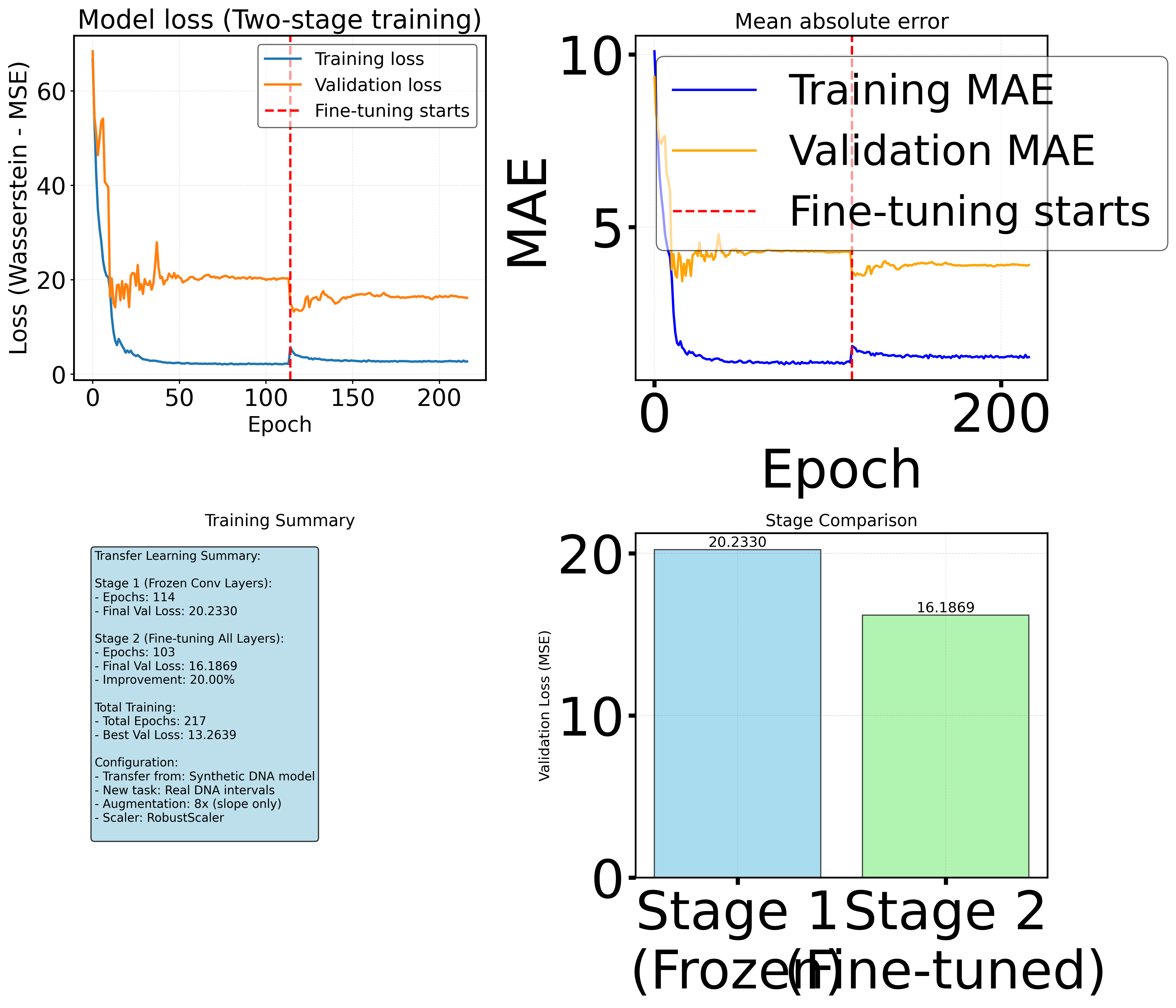

**Supplementary Figure 11**: Training and validation loss curves for the transfer learning model. The figure shows the loss history for the two-stage transfer learning model trained using the custom Wasserstein-MSE loss function. During the initial training phase, both training and validation losses decrease and converge at approximately 115 epochs, although the validation loss exhibits noisier behaviour at early epochs. Upon initiation of fine-tuning, the validation loss continues to decrease while the training loss increases slightly, consistent with transfer-learning behaviour as the model adapts to the target domain. A separation between training and validation loss is observed, which is expected given the limited size of the biological training dataset and the domain shift between synthetic and biological DNA samples. Importantly, this gap stabilises at approximately 180 epochs rather than continuing to diverge, indicating convergence of model optimisation rather than progressive overfitting.
